# The FUSED LEAVES1/*ADHERENT1* Regulatory Module Is Required For Maize Cuticle Development And Organ Separation

**DOI:** 10.1101/2020.02.11.943787

**Authors:** Xue Liu, Richard Bourgault, Josh Strable, Mary Galli, Zongliang Chen, Jiaqiang Dong, Isabel Molina, Andrea Gallavotti

**Affiliations:** Waksman Institute of Microbiology, Rutgers University, Piscataway, NJ 08854-8020, USA; Department of Biology, Algoma University, Sault Ste. Marie, ON P62A 2G4, Canada; School of Integrative Plant Science, Plant Biology Section, Cornell University, Ithaca, NY 14853, USA; Department of Plant Biology, Rutgers University, New Brunswick, NJ 08901, USA

## Abstract

In land plants all aerial epidermal cells are covered by the cuticle, an extracellular hydrophobic layer. The cuticle represents a primary barrier between cells and the external environment, provides protection against abiotic and biotic stresses, and prevents organ fusion during development. Here we report the cloning and characterization of a classic mutant of maize called *adherent1* (*ad1*), first described a century ago, and we show that *AD1* encodes a 3-KETOACYL-CoA SYNTHASE involved in the deposition of cuticular wax on the epidermis of leaves and inflorescences. *ad1* mutants show decreased amounts of various wax components as well as a range of organ fusion defects during vegetative and reproductive development. Accordingly, we find that *AD1* is strongly expressed in the epidermis of various developing organs where it is directly regulated by the MYB transcription factor FUSED LEAVES1 (FDL1), which in turn controls a series of additional genes involved in cuticle formation. Altogether, our results identify a major pathway of cuticle biosynthesis essential for the development of maize plants, and a key regulatory module that is conserved across monocot and eudicot species.

**One sentence summary:** The classic maize mutant *adherent1*, first isolated a century ago, is affected in an enzyme responsible for cuticle formation that is regulated by the MYB transcription factor FUSED LEAVES1.

## INTRODUCTION

The plant cuticle covers the aerial epidermis of all land plants, providing protection against external environmental stresses, including drought, extreme temperatures, UV light, pests and pathogens, and represents one of the most important adaptations for the terrestrial life of land plants (Yeats and Rose, 2013). It is synthesized by the plant epidermis, a single sheet of cells derived from the L1 layer of meristems that provides a necessary barrier for separating individual developing organs (Javelle et al., 2011). The cuticle structure, composition and properties influence cuticle permeability, which has been hypothesized to be critical in preventing organ fusion in early organ development (Yephremov et al., 1999; Pruitt et al., 2000; Ingram and Nawrath, 2017). Much of our understanding of postgenital fusion, defined as fusions occurring at the margins of fully formed organs (Specht and Howarth, 2015), of the epidermis comes from studies in several species including *Catharanthus roseus*, maize and Arabidopsis (Walker, 1975; Li et al., 2016; Vialette-Guiraud et al., 2016). Organ fusion has been observed in several cuticular mutants of Arabidopsis, such as *wax1*, *wax2*, *fiddlehead/kcs10* (*fdh*), *lacerata*, *hothead*, *bodyguard*, *desperado* and *long-chain acyl-CoA synthetases1 long-chain acyl-CoA synthetase2* (*lacs1* and *lacs2*) (Jenks et al., 1996; Yephremov et al., 1999; Pruitt et al., 2000; Wellesen et al., 2001; Chen et al., 2003; Kurdyukov et al., 2006b; Kurdyukov et al., 2006a; Panikashvili et al., 2007; Weng et al., 2010). In maize, mutations in *CRINKLY4* (*CR4*), a receptor kinase, affect epidermal cells and lead to adhesion events taking place between organs (Becraft et al., 1996; Becraft et al., 2001). Most of these mutations cause organ fusion defects both in juvenile vegetative and adult reproductive organs, and display cuticle defects due to mutations in genes directly involved in cutin or cuticular wax formation, the main components of the cuticle.

Cutin is a polyester matrix composed mainly of glycerol and long-chain (C16 and C18) hydroxy fatty acid monomers (Kolattukudy, 1980; Graca et al., 2002). Cuticular waxes are instead composed of a mixture of aliphatic and alicyclic compounds. Wax aliphatics are derived from very-long-chain fatty acids (VLCFAs; C20–C34) (Buschhaus and Jetter, 2011). VLCFA biosynthesis begins with de novo C16 or C18 fatty acid biosynthesis in the plastid of epidermal cells. Exported to the cytosol as acyl-CoAs, further elongation up to C34 is achieved by the FATTY ACID ELONGASE (FAE) complex at the endoplasmic reticulum (ER). The FAE complex catalyzes the two-carbon condensation from malonyl-CoA to an acyl-CoA, and consists of four enzymes, 3-KETOACYL-CoA SYNTHASE (KCS), 3-KETOACYL-CoA REDUCTASE (KCR), 3-HYDROXYACYL-CoA DEHYDRATASE (HCD) and ENOYL-CoA REDUCTASE (ECR). During the elongation process, the first step is the condensation of C2 units to acyl-CoA by KCS, the enzyme that confers substrate specificity to the complex. VLCFAs are subsequently transformed into primary alcohols and wax esters, or into aldehydes, alkanes, secondary alcohols and ketones (Kunst and Samuels, 2003; Samuels et al., 2008), and are exported to the cuticle by transporters at the plasma membrane (Pighin et al., 2004; Panikashvili et al., 2007).

Cuticular wax biosynthesis is regulated at the transcriptional level mainly by members of the MYB and AP2/ERF families of transcription factors (TFs). In Arabidopsis, WAX INDUCER1(WIN1)/SHINE1(SHN1), SHN2, SHN3 and WRINKLED4 (WRI4) are AP2/ERF TFs that directly activate genes in the wax synthesis pathway (Aharoni et al., 2004; Broun et al., 2004; Park et al., 2016), while DECREASE WAX BIOSYNTHESIS (DEWAX) and DEWAX2, negatively regulate wax production (Go et al., 2014; Kim et al., 2018). The MYB TF family also includes important regulators of cuticular wax biosynthesis, such as AtMYB16, AtMYB30, AtMYB94, AtMYB96, and AtMYB106 (Raffaele et al., 2008; Seo et al., 2011; Oshima et al., 2013; Lee and Suh, 2015). AtMYB16 and AtMYB106 were shown to regulate cuticle biosynthesis through direct activation of WIN1/SHN1 and cuticle biosynthetic genes in standard conditions (Oshima et al., 2013), while AtMYB30, AtMYB94 and AtMYB96 were found to activate wax biosynthetic genes under biotic or abiotic stress (Raffaele et al., 2008; Seo et al., 2011; Lee and Suh, 2015). Recently, in maize, two MYB family TFs, FUSED LEAVES1/MYB94 (FDL1) and GLOSSY3/MYB97, were identified as positive regulators of leaf cuticular wax accumulation. While *glossy3* mutants did not show any fusion defects, *fdl1* mutants showed organ fusion during the juvenile phase (Liu et al., 2012; La Rocca et al., 2015).

In this study, we characterized the *adherent1* (*ad1*) mutant of maize and show that *AD1* encodes a 3-KETOACYL-COA SYNTHASE required for cuticular wax biosynthesis. Furthermore, using genome-wide DNA binding analysis we show that *AD1* and a series of other enzymes involved in cuticle formation are directly activated by FDL1/MYB94. These findings establish a major pathway for cuticle formation in maize, and highlight a common module for the regulation of cuticle biosynthesis, involving MYB TFs and *KCS* genes, that is conserved between eudicot and monocot species.

## RESULTS

### The *adherent1* mutant shows abnormal fusions between organs and cells

In search of modifiers of the *ramosa1 enhancer locus2* (*rel2*) mutant, we performed an ethyl methanesulfonate (EMS) enhancer/suppressor screen of the pleiotropic *rel2-ref* mutant in the A619 genetic background (Liu et al., 2019). Among the M2 families segregating developmental defects we noticed a strong upright tassel branch phenotype segregating in one family (M2-92-224) as a single recessive locus (Supplemental Figure 1). After generating an F2 population for positional cloning purposes, we determined that this phenotype was segregating independently of the *rel2-ref* mutation and strongly resembled the phenotype of a classic maize mutant, originally isolated a century ago, called *adherent1* (*ad1*) (Kempton, 1920; Sinha and Lynch, 1998). F1 plants obtained from crosses between the mutant found in the M2-92-224 family and previously isolated alleles (*ad1-109D* and *ad1-110E*) failed to complement the upright tassel branch phenotype, indicating that the mutant identified in M2-92-224 was a new allele of *ad1*, and we therefore renamed it *ad1-224*.

Subsequently, we carefully analyzed the mutant phenotype of the *ad1-224* allele. The most notable phenotypes of *ad1* plants were the adherence of leaves in the juvenile stages of seedling development and the fusion of tassel branches in mature plants. In particular, the first, second, and occasionally third leaf blades upon germination were fused with themselves or with each other, giving rise to a characteristic phenotype in seedlings (Figure 1A). Normal maize tassels are characterized by long branches borne at the base of a long axis, the central spike (Figure 1B). Both long branches and the central spike are covered in spikelets, specific structures, enclosed by two glumes, that contain two florets. In mature *ad1* plants, all tassel branches were completely attached to each other and the central spike (Figure 1B), while spikelets appeared fused together and shriveled, with partially emerging anthers (Figure 1C,D, Supplemental Figure 1). Within each floret, anthers frequently showed fusion as well (Figure 1E).

**Figure 1.**
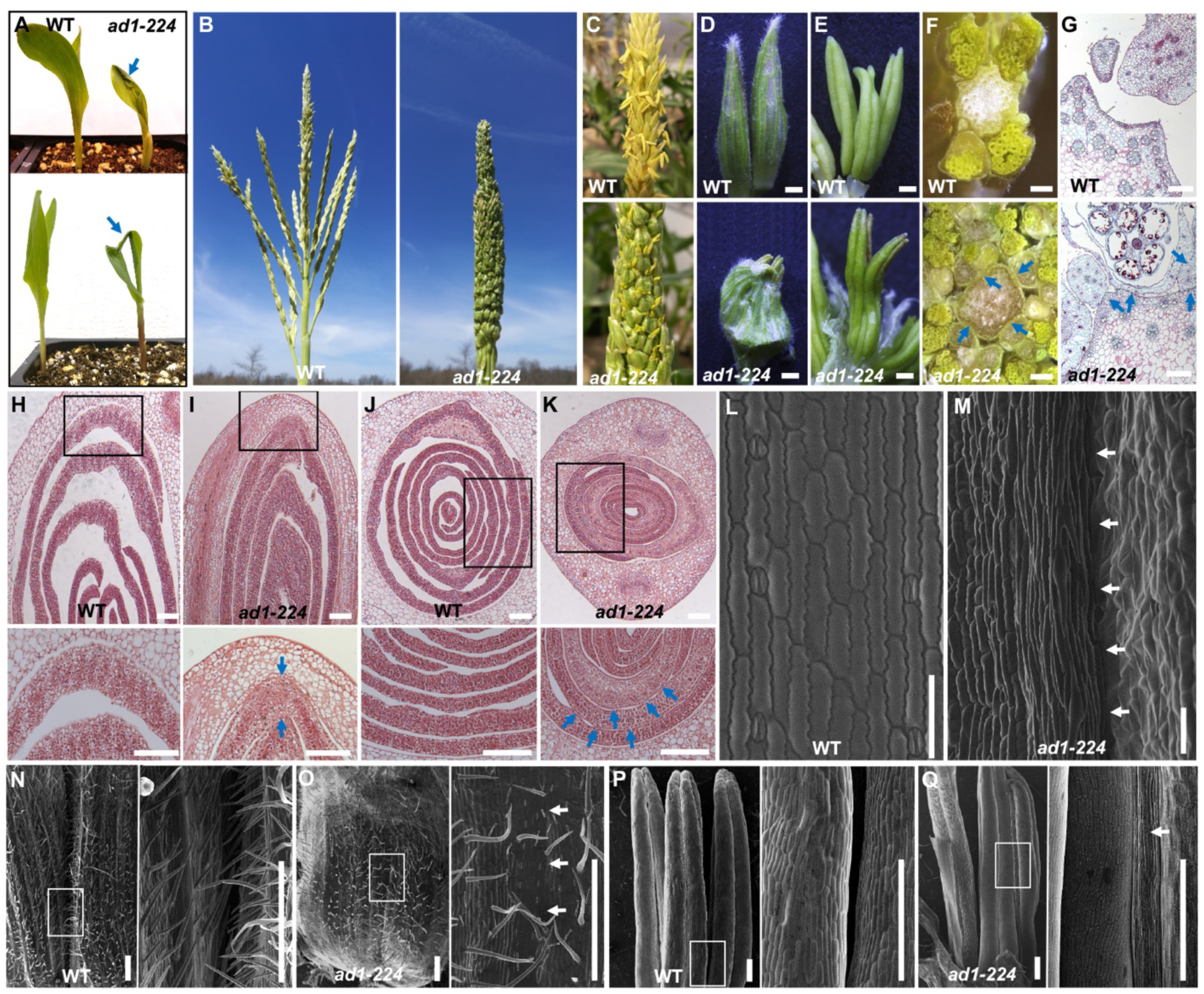
The *adherent1* mutant shows organ fusion defects. (A) The seedling phenotype of wild type and *ad1-224* plants. The second leaf is rolled and fused to the adaxial surface of first leaf (upper image). The second leaf and first leaf show fusion at the leaf tip (lower image). (B) Mature tassel phenotype. Tassel branches are fused together in *ad1-224*. (C and D) In *ad1* tassel spikelets adhere to each other, and the glumes of adjacent spikelets are fused together. Scale bars, 0.1cm. (E) Anthers in the same floret show fusion in *ad1-224*. Scale bars, 0.1cm. (F) Cross sections of wild type and *ad1-224* mature tassels. Scale bars, 0.1cm. (G) Saffranin-O Alcian Blue staining of transverse sections of wild type and *ad1-224* mature tassel. Scale bars, 0.2mm. (H-K) Longitudinal and transverse sections of wild type (H and J) and *ad1-224* (I and K) germinating seedlings. Higher magnification (lower image) of the area framed in upper image. Scale bars, 0.2 mm. (L and M) SEMs of wild type leaves (L) show no fusion defects, while a seedling leaf blade curled up and fused to itself in *ad1-224* mutants (M). Scale bars, 100 μm. Arrows point to regions of fusion. (N and O) Spikelet phenotype of wild type and *ad1-224* mutants. Note that the glumes of adjacent spikelets are separated in wild type but fused in *ad1-224* mutants. Higher magnification (right image) of the area framed in left image. (P and Q) Anthers of same floret are separated in wild type tassels, but fused in *ad1-224* mutants. Higher magnification (right image) of the area framed in left image. Scale bars in N-Q, 500 μm.

Longitudinal and transverse sections of germinating shoots, tassels and spikelets of wild type and *ad1* mutants revealed widespread organ fusion defects. For example, the coleoptile and the first leaf, the first and second leaves were fused in *ad1* germinating shoots (Figure 1H-K), with both the adaxial and abaxial sides of the blade participating in fusion events (Figure 1I,K), while *ad1* tassel branches and glumes adhered to each other and to the central spike (Figure 1F,G). Unlike tassels, mature ears of *ad1* mutants did not show a visible phenotype (Supplemental Figure 1).

To examine seedling and tassel fusion events in greater detail, we used scanning electron microscopy (SEM). *ad1* mutants displayed several types of fusion events i.e. a seedling leaf curled up and fused to itself (Figure 1M), the second leaf fused to the surface of first leaf along the margin and the macrohair from one surface fused to epidermal cells on the other surface (Supplemental Figure 1). Areas of fusions between juxtaposed epidermal surfaces were quite extensive and often appeared seamless, making it difficult to identify the boundary between the partners of the fusion event. Similar extensive and seamless fusion events of distinct epidermal cell layers were observed in *ad1* tassel glumes, which were reduced in size, and anthers (Figure 1N-Q; Supplemental Figure 1). We also checked developing ear spikelets using SEM and observed fusion events in adjacent glumes (Supplemental Figure 1). The seemingly normal appearance of mature ears therefore may be simply due to the reduced outgrowth of glumes in ear spikelets when compared to tassel spikelets. Taken together these observations indicate that the organ adherence of *ad1* mutants were caused by epidermal fusions among and between different tissues and organs in both juvenile and reproductive stages, and suggest that AD1 plays an important role in establishing epidermal properties required to maintain proper organ separation throughout maize development.

### *AD1* encodes a 3-KETOACYL-CoA SYNTHASE involved in cuticular wax biosynthesis

Using an F2 segregating population and a positional cloning approach, the *ad1-224* allele was mapped to a 10.8 Mb window on chromosome 1 between SSR markers umc1147 and umc2080 (Figure 2A). We subsequently performed Bulked Segregant Whole Genome Sequence analysis to identify point mutations in coding genes within the mapping window (Dong et al., 2019). A G to A transversion was found in the *ad1-224* mutant bulk sample within the coding region of *GRMZM2G167438/Zm00001d032728* (B73v3/v4), which introduced a premature stop-codon (W210>STOP). *GRMZM2G167438/Zm00001d032728* encodes a 3-KETOACYL-CoA SYNTHASE (KCS), a key enzyme in cuticular wax biosynthesis (Yeats and Rose, 2013). To confirm that *GRMZM2G167438/Zm00001d032728* corresponded to *AD1*, we sequenced the candidate gene in three additional alleles (*ad1-109D*, *ad1-110E* and *ad1-09-2121*). Sequencing results showed that *ad1-109D* and *ad1-110E* contained an identical and unique 1067 bp insertion in the second exon, leading to a frame shift (hereafter *ad1-109D* and *ad1-110E* are referred to as *ad1-ref*; Supplemental Figure 2), while *ad1-09-2121* carried a C to T transversion within the second exon that introduced a premature stop-codon (Figure 2A,B; Q327>STOP). Mutations in all three independent alleles were predicted to produce truncated AD1 proteins (Figure 2B). These results confirmed that knockout of *GRMZM2G167438/Zm00001d032728* caused the adherence phenotype of *ad1* mutants.

**Figure 2.**
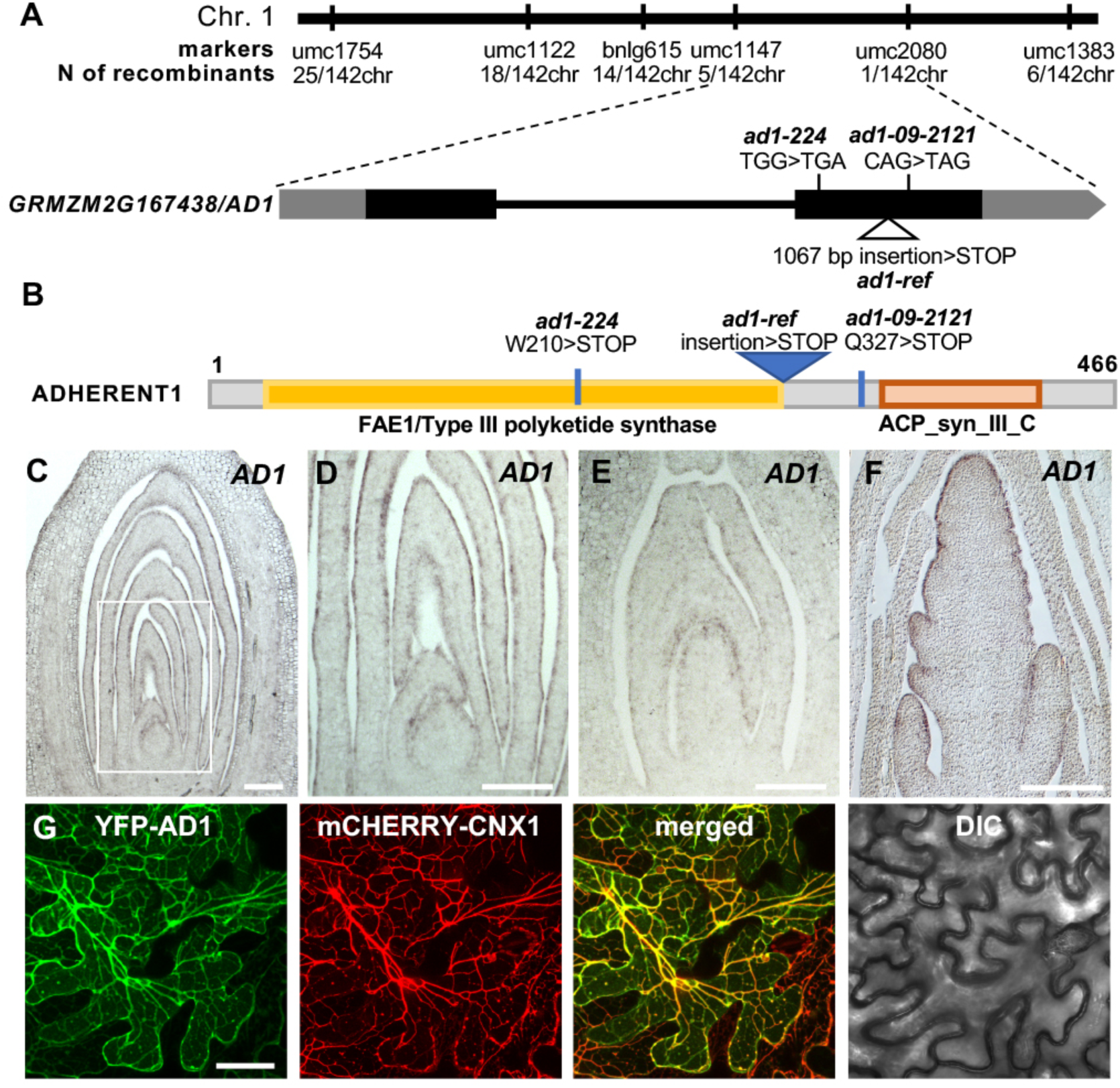
*AD1* encodes a 3-KETOACYL-CoA-SYNTHASE. (A) Positional cloning of *AD1*. Schematic representation of the *AD1* gene and the position of the mutant alleles. Exons are depicted as black rectangles and UTRs are depicted as grey rectangles. (B) Schematic representation of the AD1 protein. ACP_syn_III_C, 3-Oxoacyl-[acyl-carrier-protein (ACP)] synthase III C terminal (Pfam). (C to F) RNA *in situ* hybridizations with *AD1* antisense probes. Expression pattern in wild type germinating seedlings (C), higher magnification of the area framed in C (D), shoot region of 20 DAP embryos (E), and immature tassel (F). Scale bars, 0.2 mm. (G) Confocal images of YFP-AD1 shows co-localization with the ER marker mCHERRY-CNX1 in *N. benthamiana* leaf epidermal cells. Scale bars, 50 μm.

AD1 is an enzyme of 466 amino acids containing a conserved domain that includes substrate binding, active and product binding sites (Figure 2B). Based on qRT-PCR and available RNA-seq data, and consistent with the severe phenotype observed in seedlings and tassels, *AD1* showed high levels of expression in tassels, ears and seedling leaves, moderate levels in silks and embryos, and weak levels in endosperm, mature leaves and roots (Supplemental Figure 2). To characterize the expression pattern of *AD1* in more detail, we carried out RNA *in situ* hybridizations in germinating shoots, 20 DAP embryos and immature tassels. In all tissues tested, *AD1* showed strongest expression in the epidermal L1 layer of young leaves, embryos and tassels (Figure 2C-F). To determine its subcellular localization, we performed confocal co-imaging of a YFP-AD1 fusion protein and the endoplasmic reticulum (ER) marker mCHERRY-CNX1 (Gao et al., 2012) in *N. benthamiana* leaf epidermal cells. Strong co-localization of the YFP-AD1 and mCHERRY-CNX1 signals was observed (Figure 2G). Altogether, the expression of *AD1* in epidermal cells and the subcellular localization of AD1 in the ER are consistent with the role of AD1 in cuticular wax biosynthesis (Yeats and Rose, 2013).

The KCS family is plant specific and shows extensive genetic redundancy (Yeats and Rose, 2013). The Arabidopsis genome contains 21 *KCS* genes (Costaglioli et al., 2005; Joubes et al., 2008), and several *KCS* genes have been previously described for their role in the synthesis of VLCFAs (James et al., 1995; Todd et al., 1999; Yephremov et al., 1999; Fiebig et al., 2000; Gray et al., 2000; Pruitt et al., 2000; Franke et al., 2009; Lee et al., 2009; Kim et al., 2013). A homology-based search showed that the maize B73v3 reference genome contains 28 maize *KCS* genes. Neighbor-joining analysis classified the 21 Arabidopsis KCS proteins and 28 maize KCS proteins into several different clades, and placed AD1 within the clade that includes Arabidopsis KCS3, KCS12 and KCS19. FDH/KCS10, whose mutants also showed organ fusion phenotypes in Arabidopsis (Yephremov et al., 1999; Pruitt et al., 2000), belongs to a separate distant clade (Supplemental Figure 3). To understand why *ad1* single mutants showed a severe phenotype in maize, we analyzed the expression level of the 28 *KCS* genes in different tissues based on publicly available RNA-seq datasets (Stelpflug et al., 2016). In general, *KCS* genes were broadly expressed in all tissues throughout maize development, although some differences could be observed based on expression levels in aerial vs. root tissue (Supplemental Figure 2). Taken together, these results suggest that loss of *AD1* function cannot be completely compensated by related family members despite showing largely overlapping expression patterns with them (i.e. *GRMZM2G162508/ZmKCS15*).

### Cuticular wax biosynthesis and deposition are defective in *ad1* mutants

To investigate whether loss of AD1 function influenced cuticular wax biosynthesis and deposition, we tested the “lotus effect”, a self-cleaning mechanism of many plant leaves in which water tends to roll to the ground as droplets, collecting and washing particles and debris from the leaf surface in the process. The efficiency of this self-cleaning mechanism has been correlated with the abundance of epicuticular wax crystals (Barthlott and Neinhuis, 1997). Wild type and *ad1* mutant seedlings were misted with water, and water droplets accumulated only on the surface of *ad1* mutant leaves (Figure 3A), suggesting epicuticular wax crystal defects. We therefore analyzed the deposition of epicuticular wax crystals on the leaf surface by SEM. Strikingly, the number of epicuticular wax crystals was considerably reduced and wax crystals appeared smaller in *ad1* mutant leaves than wild type (Figure 3B). We also quantified the accumulation of toluidine blue stain in germinating coleoptiles, an assay used to measure cuticle permeability, and detected a higher accumulation of the dye in *ad1* mutants (Figure 3C,D), suggesting defects in cuticle formation during embryogenesis (Tanaka et al., 2004; Doll et al., 2020). In addition, we also examined cuticular wax accumulation by measuring cuticular transpiration and chlorophyll leaching (Figure 3E,F), which are frequently used to examine cuticular defects in leaves (Chen et al., 2003). This analysis showed that both cuticular transpiration, measured as loss of leaf weight, and chlorophyll extraction occurred more rapidly in *ad1* mutant alleles relative to wild type, further confirming the observed defects in cuticular wax accumulation of *ad1* mutants.

**Figure 3.**
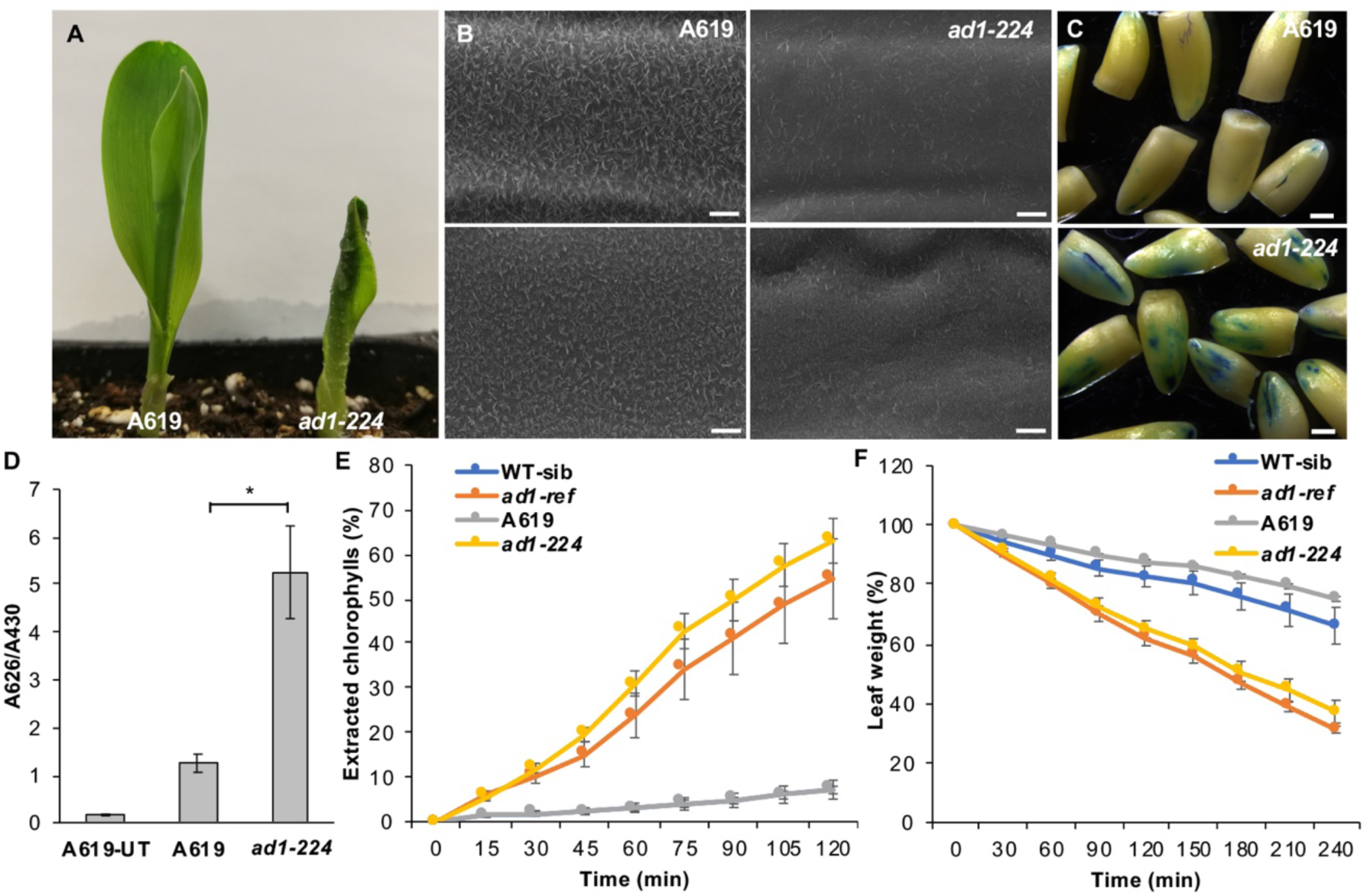
Epicuticular wax crystals and transpiration analysis. (A) Leaves of wild type and *ad1-224* seedlings misted with water. (B) SEM images of epicuticular wax crystals on the third leaf sheath (top) and the third leaf lamina (bottom) in wild type and *ad1-224* mutants. Scale bars, 5 μm. (C) Toluidine blue tests on etiolated coleoptiles. Scale bars, 1 mm. (D) Quantification of toluidine blue uptake by the coleoptile of young seedlings, normalized to chlorophyll content. n=5 (5 seedlings per repetition; * Student-test P < 0.001). Error bars represent SD. (E) Chlorophyll leaching assays. Extracted chlorophyll contents at individual time points were expressed as percentages of that at 24 h after initial immersion. Error bars show SD, n=3. (F) Water loss assays. Error bars show SD, n=3.

The reduced amount of epicuticular wax crystals on the surface of *ad1* leaves together with the water loss and chlorophyll leaching assays suggested that *AD1* is involved in cuticular wax biosynthesis. Therefore, we evaluated the amount and composition of cuticular waxes from the third leaves of wild type and *ad1* plants using gas chromatography-mass spectrometry (GC-MS) and gas chromatography coupled to a flame ionization detector (GC–FID) (Figure 4 and Supplemental Figure 4). The amount of total wax was significantly lower in *ad1-224* (∼18%) and *ad1-ref* (∼23%) than in wild type leaves (Figure 4A, Supplemental Figure 4). Maize leaf cuticular waxes are composed of a mixture of compounds including VLCFAs, alkanes, alcohols, aldehydes, ketones, wax esters and alicyclic compounds (Bourgault et al., 2020). All components were significantly decreased in *ad1-224* compared with wild type, with the exception of alkanes (Figure 4A). Similar results were obtained with the *ad1-ref* mutant, although primary alcohols and fatty acids were also not significantly different from wild type levels (Supplemental Figure 4). The primary alcohol fraction was the most abundant component class in juvenile leaf waxes (Figure 4A, Supplemental Figure 4), with the C_32:0_ primary alcohol being the dominant homolog, as expected (Bianchi et al., 1978). In fact, this single component (C_32:0_ primary alcohol) constituted over 60% of the overall wax load in the samples studied. The level of C_32:0_ primary alcohol in *ad1-224* and *ad1-ref* mutants was decreased by approximately 11% and 13% relative to the wild type, respectively (Figure 4B, Supplemental Figure 4). Another abundant component was C_32:0_ aldehyde, which constituted about 25% of the total wax load. In *ad1-224* and *ad1-ref* mutants, the C_32:0_ aldehyde load was reduced by approximately 30% and 48%, respectively, compared with values in wild type samples (Figure 4D, Supplemental Figure 4). Less abundant components, including C_30:0_-C_34:0_ fatty acids, C_30:0_ and C_34:0_ aldehydes, C_33:0_ alkane and all four identified alicyclic compounds, showed lower concentrations in *ad1-224* mutants than in control samples (Figure 4C,E,G). In *ad1-ref* mutants, leaf waxes showed reduced amounts of C_34:0_ fatty acid, C_28:0_-C_34:0_ aldehydes, C_33:0_ and C_39:0_ alkanes, tocopherolin and campesterol (Supplemental Figure 4). Interestingly, all components of the wax ester fraction, which corresponds to less than 5% of the overall wax load, were equally affected in both alleles of *ad1* (Figure 4F, Supplemental Figure 4). Altogether, *ad1* mutants had reduced amounts of cuticular wax components with acyl chains longer than 31 carbons (with the exception of the primary alcohol fraction), suggesting a key role for AD1 in the formation of wax compounds with more than 31 carbons.

**Figure 4.**
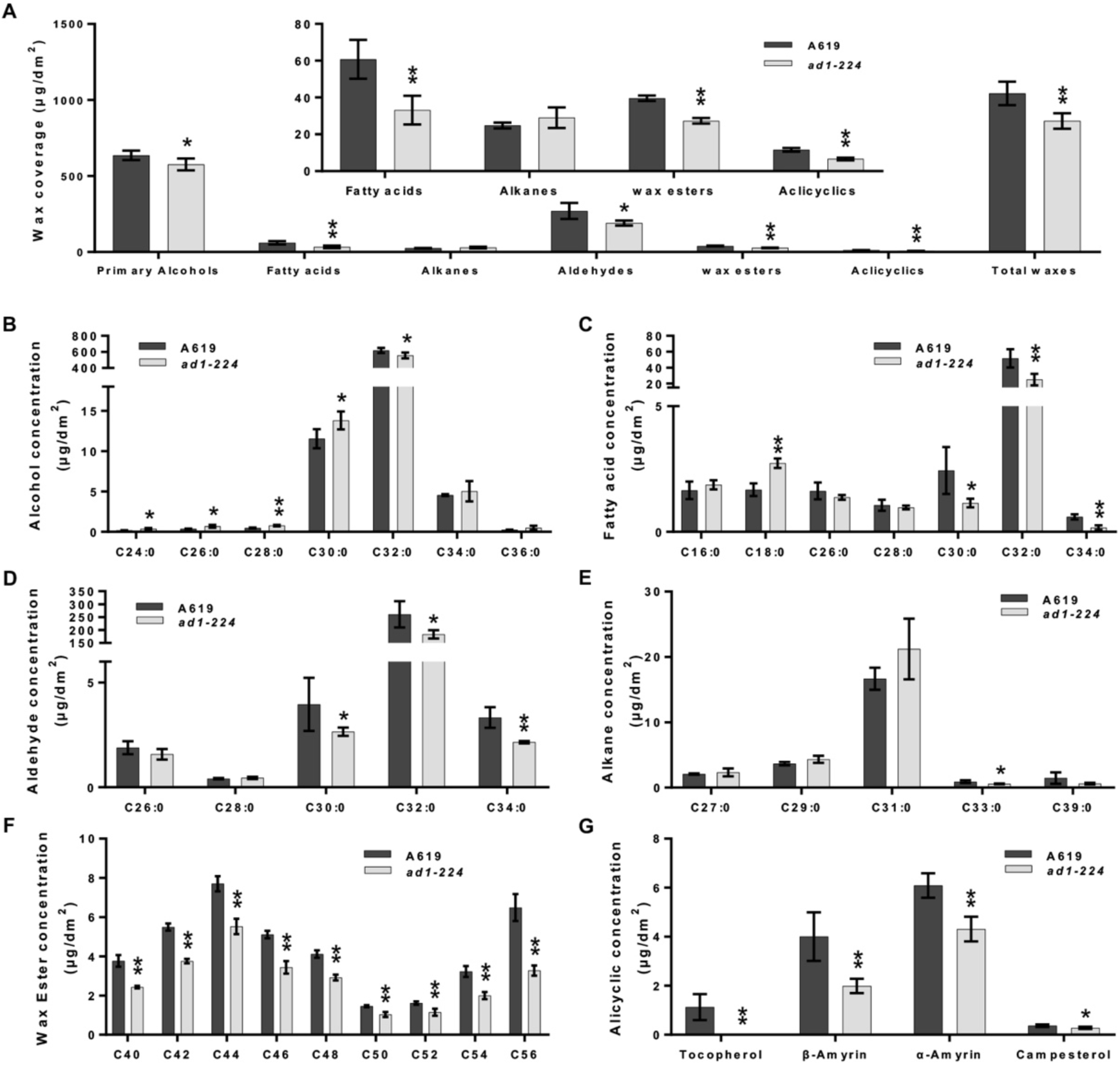
*ad1* cuticular wax analysis. (A) Total wax coverage and amount of each wax class in *ad1-224* mutants and wild type leaves. The inset shows the less abundant wax classes at a different scale to more clearly visualize significant differences. (B to G) Concentration of individual components in each wax class: primary alcohol (B), fatty acid (C), aldehyde (D), alkane (E), wax ester (F) and alicyclic (G). Means of 4 replicates and SD are reported. The asterisks represent significant difference determined by the Student’s two-tail *t* test; *, P < 0.05; **, P < 0.01.

### The MYB transcription factor FUSED LEAVES1 positively regulates *AD1* **Expression**

The previously characterized maize mutant *fused leaves1* (*fdl1*) has a phenotype similar to *ad1,* in particular regarding seedling leaf blade fusion and cuticular wax accumulation defects. *FDL1* (*GRMZM2G056407/Zm00001d022227*) encodes MYB94, a MYB transcription factor (La Rocca et al., 2015) whose Arabidopsis homologs are known to be involved in the regulation of cuticular wax biosynthesis (Raffaele et al., 2008; Seo et al., 2011; La Rocca et al., 2015). To understand whether *AD1* was regulated by FDL1/MYB94, we obtained a transposon insertion (mu1092890) in the *FDL1* gene, which disrupts its second exon (Figure 5A). Homozygous *fdl1-Mu* mutant plants showed organ fusion defects at the seedling stage (Supplemental Figure 5). Similar to *ad1* mutants, a reduced amount of epicuticular wax crystals, increased loss of epidermal water and faster chlorophyll leaching were also observed in *fdl1-Mu* plants (Supplemental Figure 5) mirroring what was previously reported for another *fdl1* allele (La Rocca et al., 2015). Using RNA *in situ* hybridizations, we determined that *FDL1* showed strong expression in the epidermal layer of young leaves and tassels, in a pattern remarkably similar to *AD1* (Figure 5B, Supplemental Figure 5). We next generated *ad1;fdl1* double mutants by crossing *ad1-ref* with *fdl1-Mu* plants. At the seedling stage, both single mutants and double mutants showed the characteristic leaf fusion phenotype (Figure 5C). SEMs of the leaf surface showed that the epicuticular wax crystal deposition defects of *ad1-ref* were enhanced by the *fdl1* mutation (Figure 5D). Accordingly, water loss by cuticular transpiration was slightly faster in *ad1-ref;fdl1-Mu* double mutants than in *ad1-ref*, *fdl1-Mu* and wild type samples, and chlorophyll leaching occurred more rapidly in *ad1-ref;fdl1-Mu* double mutants compared with each single mutant and wild type (Figure 5E,F). The accumulation of toluidine blue stain in germinating coleoptiles was also higher in *ad1;fdl1* double mutants relative to single mutants (Figure 5G,H), and in double mutant mature plants we also noticed a higher frequency of fusion events in leaves of adult plants (Supplemental Figure 6).

**Figure 5.**
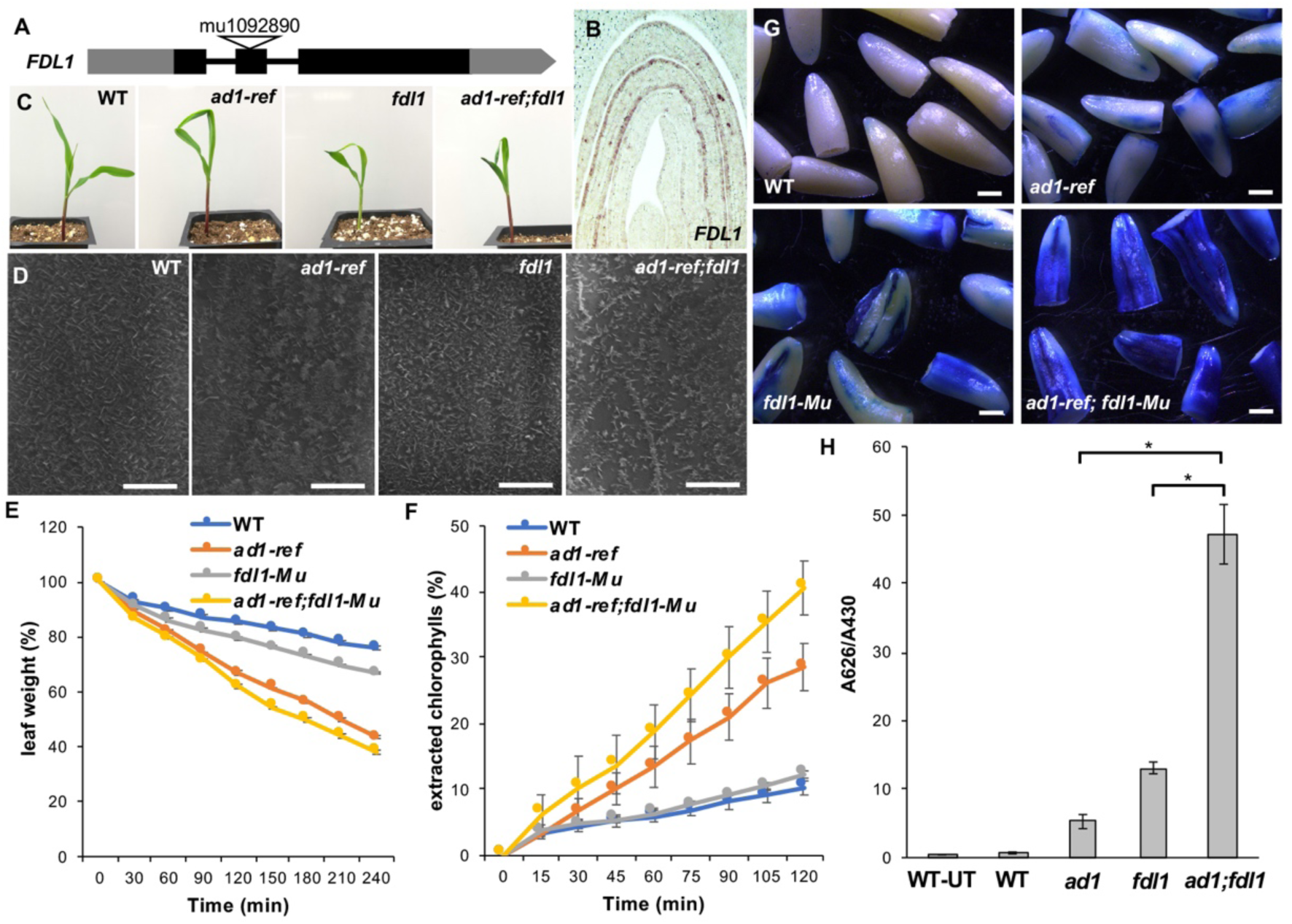
*fdl1-Mu* enhances the cuticle defects *of ad1* mutants. (A) Schematic representation of the *FDL1* gene showing the transposon insertion in the second exon (black and grey rectangles, exons and UTR regions, respectively). (B) RNA *in situ* hybridization with an *FDL1* antisense probe in germinating seedlings. (C) The seedling phenotype of wild type, *ad1-ref*, *fdl1-Mu* and *ad1-ref;fdl1-Mu*. (D) SEM images of epicuticular wax crystals on the third leaves in wild type, *ad1-ref*, *fdl1-Mu* and *ad1-ref*;*fdl1-Mu*. Scale bars, 5 μm. (E) Water loss assays of dark-acclimated plants. Error bars show SD, n=3. (F) Chlorophyll leaching assays. Extracted chlorophyll contents at individual time points were expressed as percentages of that at 24 h after initial immersion. Error bars show SD, n=3. (G) Toluidine blue assay on etiolated coleoptiles. Scale bars, 1 mm. (H) Quantification of toluidine blue uptake by the aerial parts of young seedlings, normalized to chlorophyll content (n=5, 5 seedlings per replicate; * Student-test, P < 0.001). Error bars represent SD. UT, untreated samples.

To investigate the hypothesis that FDL1/MYB94 could directly regulate the expression of *AD1* and other *KCS* genes, we performed DAP-seq, an *in vitro* DNA-TF binding assay that captures genomic DNA binding events in their native sequence context (O’Malley et al., 2016; Galli et al., 2018). *In vitro* expressed HALO-FDL1 proteins were incubated with maize genomic DNA libraries and FDL1-bound DNA fragments were identified using next-generation sequencing. In total, 10,028 FDL1/MYB94 binding events were detected of which 2,737 (27%) were located at distances up to 10Kb upstream of the transcription start site (TSS), within the gene or 3Kb downstream of the transcription termination site (TTS) (Figure 6A), putatively targeting 2,533 unique genes. FDL1 preferentially bound to sequences containing the core CCAACC motif (Figure 6B). This consensus motif was highly similar to the motif observed previously for the distantly related P1/MYB3 TF in maize (Grotewold et al., 1994; Morohashi et al., 2012) as well as the motif identified in DAP-seq experiments for AtMYB94 and AtMYB96 (O’Malley et al., 2016). It also resembled that reported in the promoters of several cuticle-related target genes of AtMYB94 and AtMYB96, although several nucleotides outside of the core AAC motif were divergent in these genes (Seo et al., 2011; Lee and Suh, 2015).

**Figure 6.**
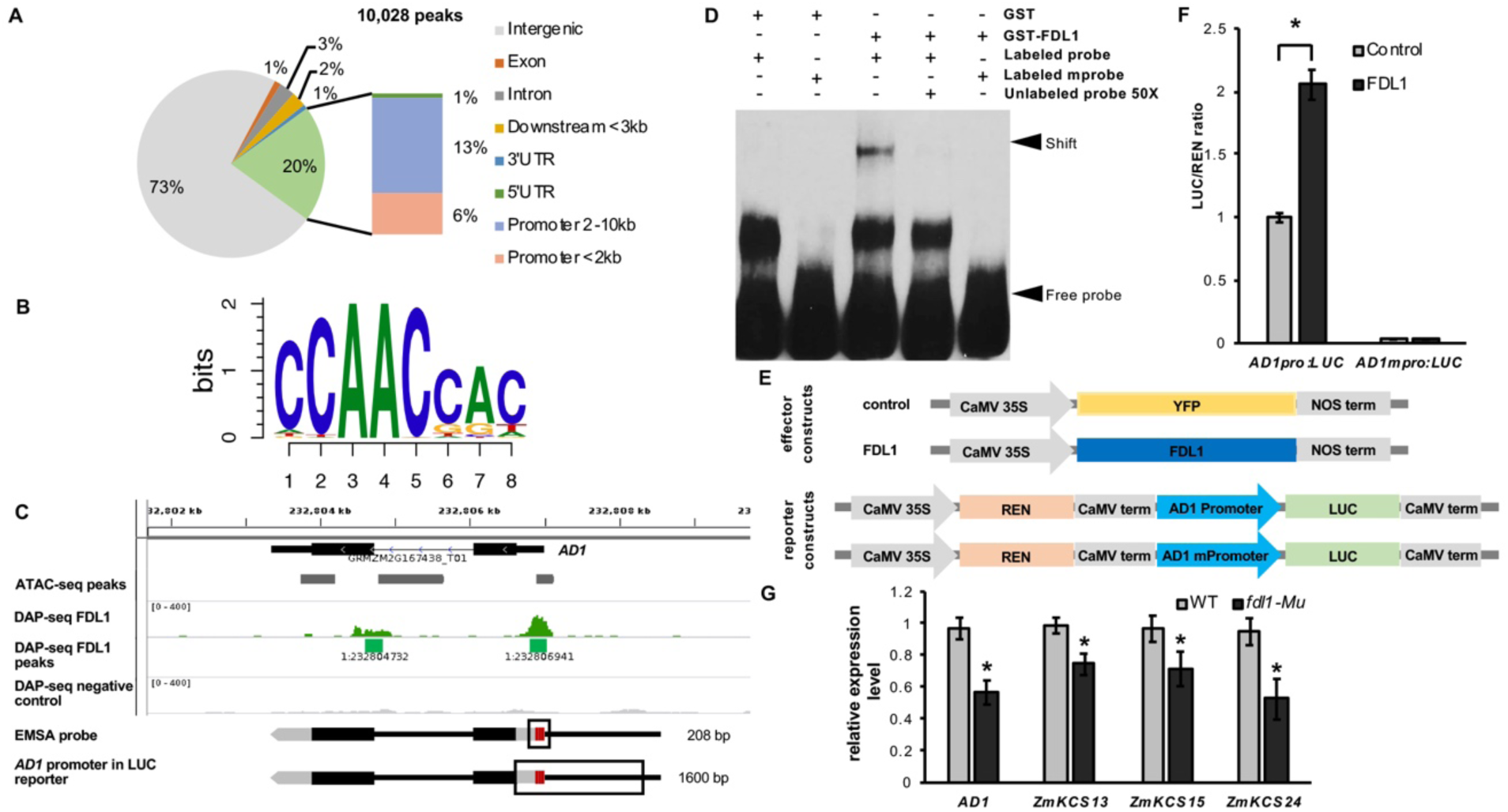
FDL1 directly regulates *AD1* expression. (A) Distribution of FDL1 peaks (qvalue e-5 Q30) within gene features. (B) Top FDL1 binding motif identified by DAP-seq. (C) FDL1 directly binds to *AD1* 5’UTR in DAP-seq. Screenshot of aligned reads. In green are significant peaks. Gene models show location of EMSA probe and the promoter used for the Luciferase assay (boxed) and the core binding motif predicted by DAP-seq marked in red lines that were mutated to generate mprobe (in EMSAs) and mPromoter (mpro, in transcriptional activation assay). (D) EMSAs show that GST-FDL1 directly bind to the promoter of *AD1*. (E) Schematic diagram of effector and reporter constructs used for transcriptional activation assay. (F) *AD1* transcriptional activation by FDL1. Error bars show SD, n=4. (G) Comparison of *AD1* and selected KCS genes expression levels between wild type and *fdl1-Mu*. The maize *UBIQUITIN* gene was used as reference. Values are means ± SD (n=3). Asterisks represent significant difference determined by the Student’s two-tail *t* test at P < 0.001.

A major functional category of genes directly bound by FDL1 included those encoding a subset of wax biosynthetic enzymes, such as homologs of Arabidopsis *KCR1*, *CER1* and *CER3*, as well as many *KCS* family genes (Supplemental Table 1)(Yeats and Rose, 2013). Overall, of the 28 *KCS* genes in maize, we observed strong peaks in putative regulatory regions of five *KCS* genes including *AD1* (Figure 6C; Supplemental Figure 7, Supplemental Table 1), suggesting that FDL1 may regulate cuticular wax biosynthesis in maize via transcriptional control of multiple *KCS* genes. An additional seven *KCS* genes also showed weak binding (adjusted p-value cutoff 0.01). Maize homologs of genes encoding the putative wax transporter AtLTPG1 and regulators of cuticular wax biosynthesis AtMYB30, AtMYB106 were also directly bound by FDL1, suggesting that cuticular wax biosynthesis and transport were broadly regulated by FDL1 (Supplemental Table 1). Many additional genes for which no direct role in wax formation has yet been described, were also found to be strongly bound by FDL1. These included *ZmLTP2/GRMZM2G010868*, encoding a lipid transfer protein whose closest Arabidopsis homolog is expressed in the L1 layer; *ZmPSS1* (*phosphatidylserine synthase1/GRMZM2G039385*), a gene involved in lipid metabolism that affects Arabidopsis pollen and meristem development (Liu et al., 2013); and *ZmEXPA1/GRMZM2G339122*, encoding alpha expansin1, a cell wall modifying enzyme (Ramakrishna et al., 2019).

As mentioned above, *AD1* was among the list of *KCS* genes directly bound by FDL1. A strong peak was located in the 5’UTR region along with a minor peak in the first intron (Figure 6C). We subsequently checked the sequence of the proximal *AD1* promoter and found that three core CCAACC motifs were present in the FDL1 binding peak region (Figure 6C, Supplemental Figure 7). Electrophoretic mobility shift assays (EMSAs) using a glutathione S-transferase (GST)-FDL1 fusion protein produced in *E. coli* were performed to test FDL1 binding to the *AD1* promoter. A 208 bp *AD1* promoter sequence containing the three core CCAACC motifs was used as the target probe with a mutated core motif sequence containing TTTTTT as a negative control. As shown in Figure 6D, GST-FDL1 bound the probe containing the CCAACC motifs resulting in a visible shift of the protein-DNA complex. This shift was not observed when GST-FDL1 was incubated with the negative control probe containing the mutated core motif. Furthermore, addition of excess amounts of unlabeled wild type probe outcompeted labeled wild type probe, indicating a specific interaction. These results indicated specific binding of FDL1 to the *AD1* promoter sequence *in vitro*. To investigate whether FDL1 binding to the *AD1* promoter could affect its transcription, we used a transient dual-luciferase (LUC) expression assay in maize protoplasts. An FDL1 effector construct and a luciferase reporter construct driven by a 1.6-kb wild type or mutated core CCAACC motif promoters of *AD1* were transiently co-expressed in protoplasts, and a 2-fold increase in the luminescence intensity compared with the control was observed (Figure 6E,F). Considerably reduced expression was observed in the reporter line driven by the mutated promoter (Figure 6F). These results indicate that FDL1 recognized the *AD1* proximal promoter region and activated its expression in maize protoplasts. In agreement with these findings, the expression level of *AD1* and other *KCS* genes (i.e. *ZmKCS13*, *ZmKCS15* and *ZmKCS24*), assessed by qRT-PCR in *fdl1* mutant embryos, were significantly reduced compared to wild type (Figure 6G), confirming that FDL1 regulates *AD1* and *KCS* transcription. These results suggest that FDL1 directly activates *AD1* expression *in vivo*, most likely by binding to the CCAACC motifs in its 5’UTR region. Collectively, these observations suggest that FDL1 functions as a positive regulator of *AD1* as well as other enzymes involved in cuticular wax biosynthesis in maize.

## DISCUSSION

Maize *ad1* mutants showed fusions between cells and organs, and fusion events were evident from embryo through tassel and floret development. These results highlight the importance of proper cuticle formation for maize shoot development, and show that AD1, a 3-KETOACYL-CoA SYNTHASE involved in cuticular wax biosynthesis, is necessary for keeping organs separated and prevent postgenital fusion events. Previously reported maize *glossy* mutants have wax accumulation defects, but unlike *ad1*, they do not display organ fusions (Bianchi et al., 1978; Moose and Sisco, 1996; Xu et al., 1997; Sturaro et al., 2005; Li et al., 2014). A similar situation is seen in Arabidopsis, where the *fdh* mutant, defective for a KCS enzyme distinct from AD1, shows severe organ fusions (Yephremov et al., 1999; Pruitt et al., 2000), while other mutations in *KCS* genes do not display similar defects (James et al., 1995; Todd et al., 1999; Yephremov et al., 1999; Fiebig et al., 2000; Gray et al., 2000; Pruitt et al., 2000; Franke et al., 2009; Lee et al., 2009; Kim et al., 2013). While redundancy in the KCS family is thought to be advantageous among higher terrestrial plant species, it is likely that AD1 and FDH, which belong to distinct clades (Supplemental Figure 3), have evolved specialized and predominant roles that cannot be complemented by other closely related family members. In addition, other gene families involved in cuticular wax as well as cutin biosynthesis when mutated also exhibit organ fusion (i.e. *wax1*, *wax2*, *lcr*, *hth* and *lacs1 lacs2*) indicating that cuticle formation involves a complex array of components, each critical for plant development (Jenks et al., 1996; Wellesen et al., 2001; Chen et al., 2003; Kurdyukov et al., 2006b; Kurdyukov et al., 2006a; Bird et al., 2007; Panikashvili et al., 2007; Weng et al., 2010). While the mechanism of how wax and cutin biosynthetic pathway mutants affect cuticle structure, composition and properties is not well understood, it is clear that proper cuticle formation which starts from L1 cells during embryo development, plays an essential role in maintaining organ separation and preventing postgenital fusions (Yephremov et al., 1999; Pruitt et al., 2000; Ingram and Nawrath, 2017). Detailed characterization of various organ fusion mutants in early stages of development should improve understanding of the molecular and cellular mechanics of how the cuticle suppresses epidermal fusion in plants. For example, recent work in Arabidopsis revealed mechanistic details of the early steps of cuticle deposition by the action of an endosperm localized subtilase and peptide/receptor complexes derived from the embryo (Doll et al., 2020).

Genome-wide DAP-seq analysis indicated that FDL1, a previously described MYB TF (La Rocca et al., 2015), directly binds to many genes involved in wax biosynthesis, including those encoding enzymes, transporters and upstream transcriptional regulators. A major category of targets bound by FDL1 and validated by qRT-PCR belongs to the *KCS* gene family, including *AD1* itself. These data suggest that FDL1 may regulate cuticular wax biosynthesis through transcriptional control of a suite of *KCS* genes. In agreement with these findings, genetic analysis showed that *ad1;fdl1* double mutants enhanced the cuticle defects of single *ad1* mutants, suggesting that other genes (i.e *ZmKCS13,15* and *24*) were the targets of FDL1 activity during juvenile development. Accordingly, both *AD1* and *FDL1* genes were predominantly expressed in epidermal cells of germinating seedlings. These data indicate that FDL1 is likely a key regulator of cuticular wax biosynthesis, and AD1 and FDL1 function is essential to form a proper barrier to prevent epidermal fusions in the early stages of maize development. Many additional genes whose role in cuticle development has yet to be described were also directly bound by FDL1, providing new avenues for the discovery of new players involved in this process.

In Arabidopsis, several MYB TFs are known to play a role in cuticle formation i.e. AtMYB16, AtMYB30, AtMYB94, AtMYB96, and AtMYB106 (Raffaele et al., 2008; Seo et al., 2011; Oshima et al., 2013; Lee and Suh, 2015). Of these, AtMYB94 and AtMYB96, the MYBs most-closely related to FDL1/ZmMYB94, have been shown by microarray, EMSA, and ChIP-PCR to directly regulate a handful of distinct cuticle biosynthetic genes (Seo et al., 2011; Oshima et al., 2013; Lee and Suh, 2015). The maize orthologs of several of these genes were also directly bound by FDL1 in our DAP-seq dataset (Supplemental Table 1). To the best of our knowledge, no regulatory information has been reported for any Arabidopsis *KCS* genes belonging to the AD1 clade, suggesting possible regulatory variation among these two species. Therefore, while a core regulatory module composed of MYB TFs and *KCS* genes appears conserved between Arabidopsis and maize, many species-specific features are likely to exist highlighting the importance of obtaining empirical data from different species.

Similar to *ad1* mutants, *fdl1* mutant plants show cuticular wax deposition defects and fusions in leaves but, unlike *ad1*, no fusion events are observed during inflorescence development. We therefore hypothesize that other closely related maize MYB family members may function redundantly to FDL1 during reproductive development. It is important to note that the Arabidopsis MYBs most closely related to FDL1 (AtMYB30/94/96), are induced by stress i.e. drought or pathogens, and positively regulate cuticular wax biosynthesis (Raffaele et al., 2008; Seo et al., 2011; Lee and Suh, 2015). These similarities suggest that the maize FDL1/*AD1* regulatory module may also enhance stress resistance by activating cuticular wax biosynthesis. Overall, these findings discovered an essential pathway for cuticle and organ development in maize, an important step to devise strategies for improving environmental adaptability and stress tolerance in variable environments for maize and other crops.

## METHODS

### Plant materials and phenotyping

The *ad1-224* allele, originally identified in an EMS mutagenesis screen for *rel2* enhancers/suppressors in the A619 background, was back-crossed to A619 before phenotypic measurements. The *ad1-ref* allele was obtained from the Maize Genetics Cooperation Stock Center (stock *ad1-109D* and *ad1-110E*), while the *ad1-9.2121* allele was originally obtained in an EMS enhancer screen of a weak *ramosa1* mutant. The *FDL1* insertion line mu1092890 (stock UFMu-13110) was obtained from the Maize Genetics Cooperation Stock Center. The phenotype of single and double mutant plants was analyzed at Rutgers University, NJ, in field and greenhouse grown plants. Student’s *t*-test was used to determine statistical significance for all measurements.

### Positional cloning

For map-based cloning of the *AD1* locus, we constructed a fine-mapping F_2_ population by crossing *ad1-224* mutant from the original M2 genetic background to the B73 inbred line. Using the PCR-based molecular markers, we mapped *ad1* to a 10.8 Mb window on chromosome 1 between markers UMC1147 and UMC2080. For Bulked Segregant Whole Genome Sequence analysis, a single bulk genomic DNA sample was obtained from 10 *ad1* plants. Library preparation and sequencing of the sample was performed by Macrogen. 150bp pair end sequencing was done on a NovaSeq 6000 Illumina instrument. The resulting fastq files were used for analysis by a previously published pipeline (Dong et al., 2019), using B73v3 as a reference genome. Two index plots were produced by the ggplot2 package in the R programming language, one for all SNPs and one for all INDELs identified by the pipelines (Dong et al., 2019). Then checked SNPs within the 10.8 Mb *ad1* mapping window using the Integrative Genomics Viewer (IGV; http://www.broadinstitute.org/igv/). The sequence of *AD1* was deposited in GenBank (accession no. MN872839) and corresponds to GRMZM2G167438_T01/Zm00001d032728_T001 (B73v3/v4).

### Microscopy

For light microscopy, thin sections (8 µm thick), were stained with Safranin O/Alcian Blue and images were acquired using a Leica DM5500B microscope equipped with a DFC450 C digital camera. For the analysis of organ fusion defects and epicuticular wax crystals, fresh tissue was imaged using a JMC-6000PLUS Scanning Electron Microscope.

For confocal microscopy, YFP-AD1 and mCHERRY-CNX1 were transiently expressed in *N. benthamiana* using Agrobacterium-mediated leaf injections. Images were obtained on a Leica SP5 confocal microscope. The YFP signal was imaged using 514 excitation and 520–575 emission, while mCHERRY was imaged using 594 excitation and 625-655 emission settings. Acquired images were analyzed using FIJI.

### Expression analysis

For quantitative real-time PCR (qRT-PCR), total RNA was extracted from different tissues using the RNeasy Plant Mini Kit (Qiagen). Retrotrascription was performed using the qScript cDNA Synthesis kit. cDNA was amplified with PerfeCTa® SYBR® Green FastMix® (Quanta Biosciences) on an Illumina Eco Real-Time PCR System. Relative expression levels were calculated using 2^−ΔΔCt^ method with *UBIQUITIN* as control. The primers used for qRT-PCR are listed in SI Appendix Table S2.

For *in situ* hybridizations, 0.2 cm B73 germinating seedlings and tassels were dissected and fixed using paraformaldehyde acetic acid (PFA). Samples were dehydrated and embedded in paraplast. Hybridizations were performed at 59°C, overnight. After several washes, samples were treated with anti-digoxygenin (DIG) antibody (Roche) and signals were detected using NBT/BCIP (Promega). The *AD1* and *FDL1* antisense probes were synthesized using T7 RNA polymerase (Promega) of *AD1* 3’UTR and *FDL1* 3’UTR cloned into pENTR223.1-Sfi and digested with EcoRI.

### Phylogenetic analysis

Full length amino acid sequences of the KCS family were obtained from Phytozome 12.1. Sequences of maize and Arabidopsis were aligned with Clustal Omega. The alignment file was then used to generate a neighbor-joining rooted tree with MEGA 5.0, applying the Bootstrap method and 1000 bootstrap replications (Simmons and Freudenstein, 2011).

### Water loss and chlorophyll leaching assays

For water loss assays, 3-week-old plants grown in greenhouse potting media were used. Whole shoots of dark-acclimated seedlings were excised and leaves soaked in water for 60 min in the dark. Subsequently, excess water was removed from the leaves, and 3cm pieces of leaves were weighed at the indicated time points using a precision balance. Three measurements were averaged per each time point in each genotype.

Chlorophyll leaching assays were performed on leaves of 3-week-old plants. Two grams of each leaf sample was incubated on ice for 30 min and immersed in 30 mL of 80% ethanol in 50 mL conical tubes at room temperature. Aliquots of 100 μL were removed from the solution at every 15 min after initial immersion. The amount of extracted chlorophylls was quantified by measuring absorbance at 647 and 664 nm using a spectrophotometer (Thermo Scientific). Three measurements were averaged for each time point in each genotype.

### Toluidine blue permeability test

For assessing cuticle integrity, toluidine blue tests were performed, as previously described (Tanaka et al., 2004; Doll et al., 2020). Two-day old etiolated coleoptiles were stained for 5 minutes in a toluidine blue solution (0.05% w/v) with Tween 20 (0.1% v/v) and washed in tap water. Pictures were acquired using a Leica DM5500B microscope equipped with a DFC450 C digital camera. For quantification, 5 coleoptiles were excised, placed in tubes containing 1 mL of 80% ethanol, and incubated for 4 hours in the dark until all dye and chlorophyll had been extracted. Toluidine blue absorbance of the resulting solution was analyzed using a spectrophotometer. Per each treatment, 5 repeats were performed.

### Cuticular wax analysis

Waxes were extracted from an expanded portion of juvenile leaves (3^rd^ leaf). A small section of each leaf used for the analysis was preliminary scanned by SEM to verify loss of crystal waxes in the mutants. Wax components were identified by GC-MS and quantified by gas chromatograph coupled to a flame ionization detector (GC–FID) following the protocol described in Bourgault et al (2020) (Bourgault et al., 2020).

### DAP-seq, read mapping, filtering, and peak calling

HALO-FDL1 and HALO-GST (negative control) DAP-seq samples were performed using protein expressed in the TNT rabbit reticulocyte expression system (Promega) as previously described (Bartlett et al., 2017) with the exception that 1 microgram of genomic maize B73 DNA library was added to the protein bound beads. Protein bound beads and gDNA were rotated for 1h at RT. Beads were washed four times in 1X PBS+NP40, followed by two washes with 1X PBS. Beads were transferred to a new tube and DNA was recovered by resuspending in 25μl EB and boiling for 10min at 98°C. Eluted samples were enriched and tagged with dual indexed multiplexing barcodes by performing 20 cycles of PCR in a 50μl reaction. Samples were pooled and sequenced on a NExtSeq500 with 75bp single end reads. A total of 10–30 million reads were obtained for each sample.

Reads were mapped to the B73v3 genome as described in Galli et al., 2018 (Galli et al., 2018). In brief, fastq files were trimmed using trimmomatic with the following parameters ILLUMINACLIP:TruSeq3-SE:2:30:10 LEADING:3 TRAILING:3 SLIDINGWINDOW:4:15 MINLEN:50. Trimmed reads were mapped to the B73v3 reference genome (nuclear chromosomes only) using bowtie2 v2.2.8. Mapped reads were filtered for reads containing >MA using samtools (samtools view –b –q 30) in order to restrict the number of reads mapping to multiple locations in the genome. MAPQ filtered reads were used for all subsequent analysis. Peaks were called using GEM v2.5 using the GST-HALO negative control sample for background subtraction and an FDR of 0.00001 (--q 5). Peak calling was performed with the following parameters: --d Read_Distribution_default.txt --k_min 6 --k_max 20. For visualization in the Integrative Genome Browser (IGV), bam files containing mapped reads were converted to bigwig files using deep Tools v2.5.3 bam Coverage with 10bp bin size and FPKM normalization. Peaks were associated with their closest putative target genes using ChIPseeker (Yu et al., 2015). The FDL1 DAP-seq dataset was deposited in the Gene Expression Omnibus (GEO) database (accession no. GSE142847).

### Electrophoretic Mobility Shift Assays

For EMSAs, a selected region −411 to −233 relative to *AD1* start codon (+1) within the *AD1* promoter that was enriched for core CCAACC elements was PCR amplified, gel purified, and biotinylated using the Biotin 3′ End DNA Labeling Kit (Thermo Scientific) according to the manufacturer’s recommendations. DNA binding assays were performed using the Lightshift Chemiluminescent EMSA kit (Thermo Scientific) as follows: binding reactions containing 1× Binding Buffer, 50 ng/μL polydI/dC, 2.5% glycerol, 2 μL biotinylated probe, and 1 μL purified GST-FDL1 protein were incubated at room temperature for 20 min and loaded on a 6% DNA retardation gel (Life Technologies) before transfer to a nylon membrane. Subsequent detection was carried out according to the manufacturer’s recommendations. Competition with unlabeled probe was carried out using 50-fold excess of unlabeled probe. Mutated probe was generated by annealing two complementary oligos in which the core CCAACC elements were changed to TTTTTT. Primers used are listed in Supplemental Table 2.

### Transient Transcription Dual-LUC Assay

The isolation of B73 maize mesophyll protoplasts, PEG-calcium transfection of plasmid DNA, and protoplast culture were performed as described previously (Sheen, 2001). The effector vector pRI101 was used for the expression of the *FDL1* gene driven by a 35S promoter. The reporter vector pGreenII 0800-LUC was used for detecting the transactivation activity of the *AD1* promoter. The ratio of LUC/REN activity was measured using the DLR assay system (Promega). Primers are listed in SI Appendix Table S2.

## AUTHOR CONTRIBUTIONS

X.L., J.S. and A.G. conceived the study; X.L., J.S., R.B., Z.C., M.G., J.D. and A.G. performed all experiments; X.L., R.B., M.G., J.D., I.M. and A.G. analyzed data; X.L. and A.G. wrote the manuscript with inputs from all authors.

## DATA DEPOSITION

The sequence of *AD1* was deposited in GenBank (accession no. MN872839) and corresponds to GRMZM2G167438_T01/Zm00001d032728_T001 (B73v3/v4). The FDL1 DAP-seq dataset was deposited in the Gene Expression Omnibus (GEO) database (accession no. GSE142847).

## ACKNOWLEDGMENTS

The authors wish to thank the Maize Genetics Cooperation Stock Center for seeds; Mariusz Roszkowski and Jonathan Kunkel-Jure for help with positional cloning; Erik Vollbrecht for providing the *ad1-9.2121* allele. This work was supported by the National Science Foundation (IOS#1456950) to A.G., by a National Science Foundation Postdoctoral Research Fellowship in Biology grant (IOS#1710973) to J.S., and by funding from the Canada Research Chairs program and the Natural Sciences and Engineering Research Council of Canada (RGPIN-2019-05923) to I.M.

