## Supplemental materials for "The FUSED LEAVES1/*ADHERENT1* Regulatory Module Is Required For Maize Cuticle Development And Organ Separation"

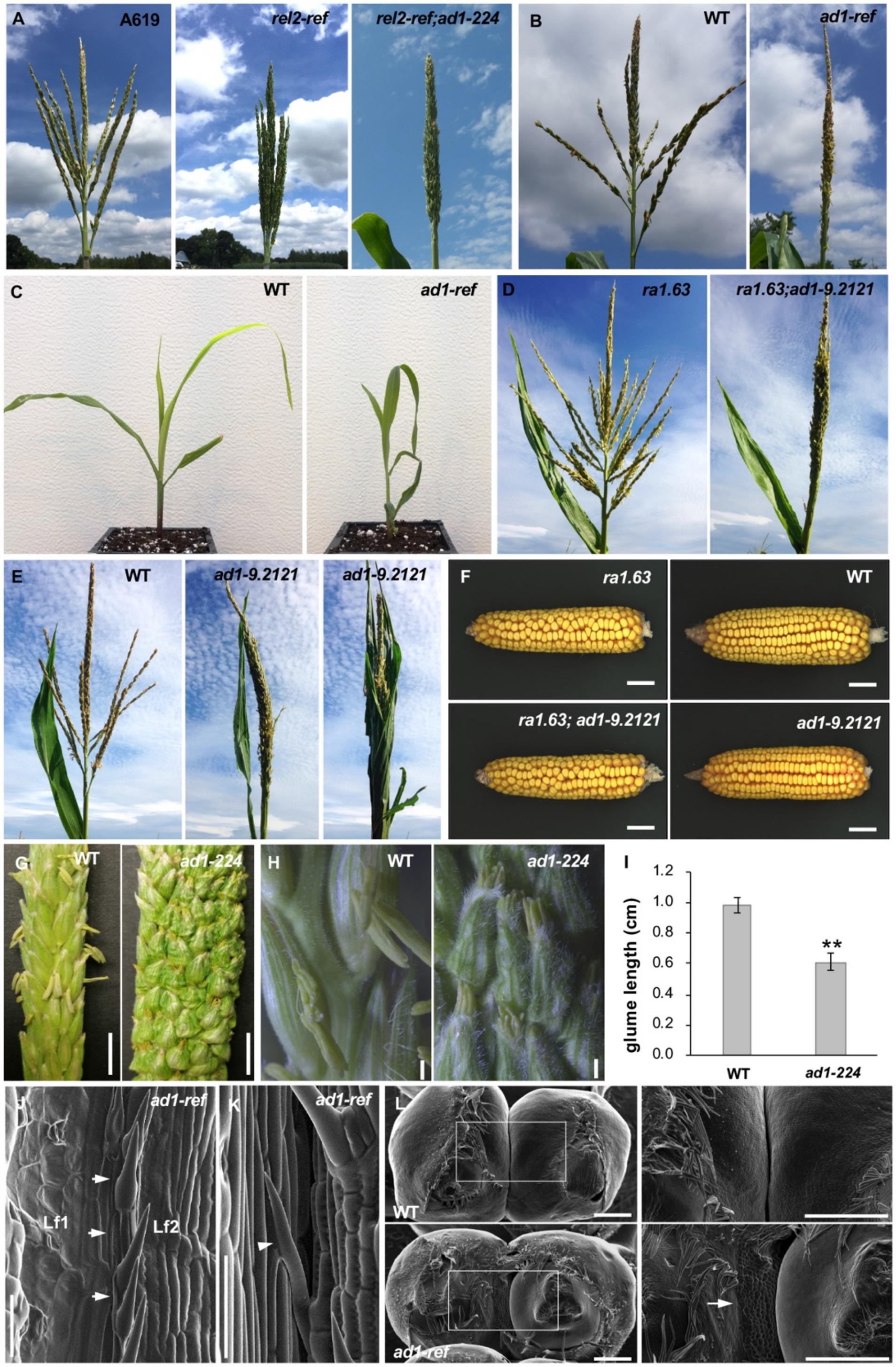


**Supplemental Figure 1.** Phenotypic characterization of *ad1* alleles.
(A) The tassel phenotype of A619, *rel2-ref* and *rel2-ref;ad1-224* original M2 family. (B) The tassel phenotype of wild type sibling and *ad1-ref* mutants. (C) The seedling phenotype of wild type sibling and *ad1-ref* mutants. (D) The original *ad1-9.2121* allele in *ra1-63.3359* Mo17 background. (E) The *ad1-9.2121* allele in BC3(B73). (F) Mature ear phenotype of *ad1-9.2121* mutants. No visible defects is observed. (G) Tassel spikelets

of wild type and *ad1-224* mutants. Scale bars, 1 cm. (H) Images of glumes in wild type and *ad1-224* mutant tassels. Scale bars, 0.1 cm. (I) Quantification of the glume length of wild type and *ad1-224* mutants (n≥20). Error bars show SD; **, $p<0.001$. (J) SEM image showing the second leaf (Lf2) fused to the surface of first leaf (Lf1) along the edge in *ad1-224* mutants. (K) SEM showing a macrohair fused to epidermal cells in *ad1-224* mutants. Scale bars in J and K, 100 µm. (L) SEM images of immature ears in wild type and *ad1-ref*. The glume of adjacent spikelets are fused in *ad1-ref* mutants. Right image is higher magnification of the area framed in left panel. Scale bars, 500 µm. Arrows point to regions of fusion events.

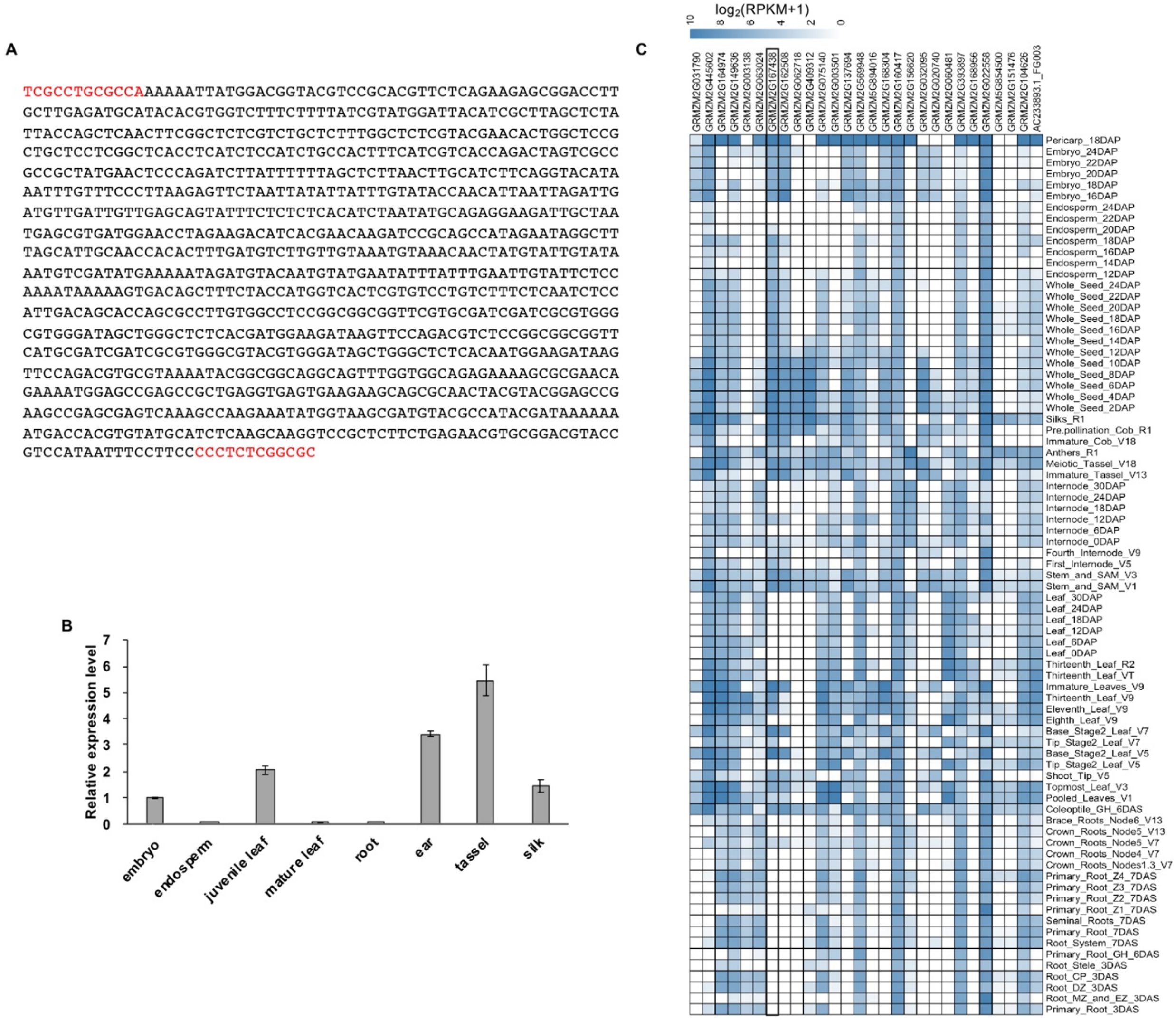


**Supplemental Figure 2.** Expression analysis of the *KCS* gene family in maize. (A) Sequence of the insertion in the *ad1-ref* allele at position +947. In red, partial coding sequence. (B) Quantitative RT-PCR of *AD1* in different maize tissues. The y axis shows the fold change relative to embryo expression levels. Error bars show SD, n=3. (C) RNA-seq expression levels of *KCS* genes in different tissues, from Stelpflug et al. 2016 (49).

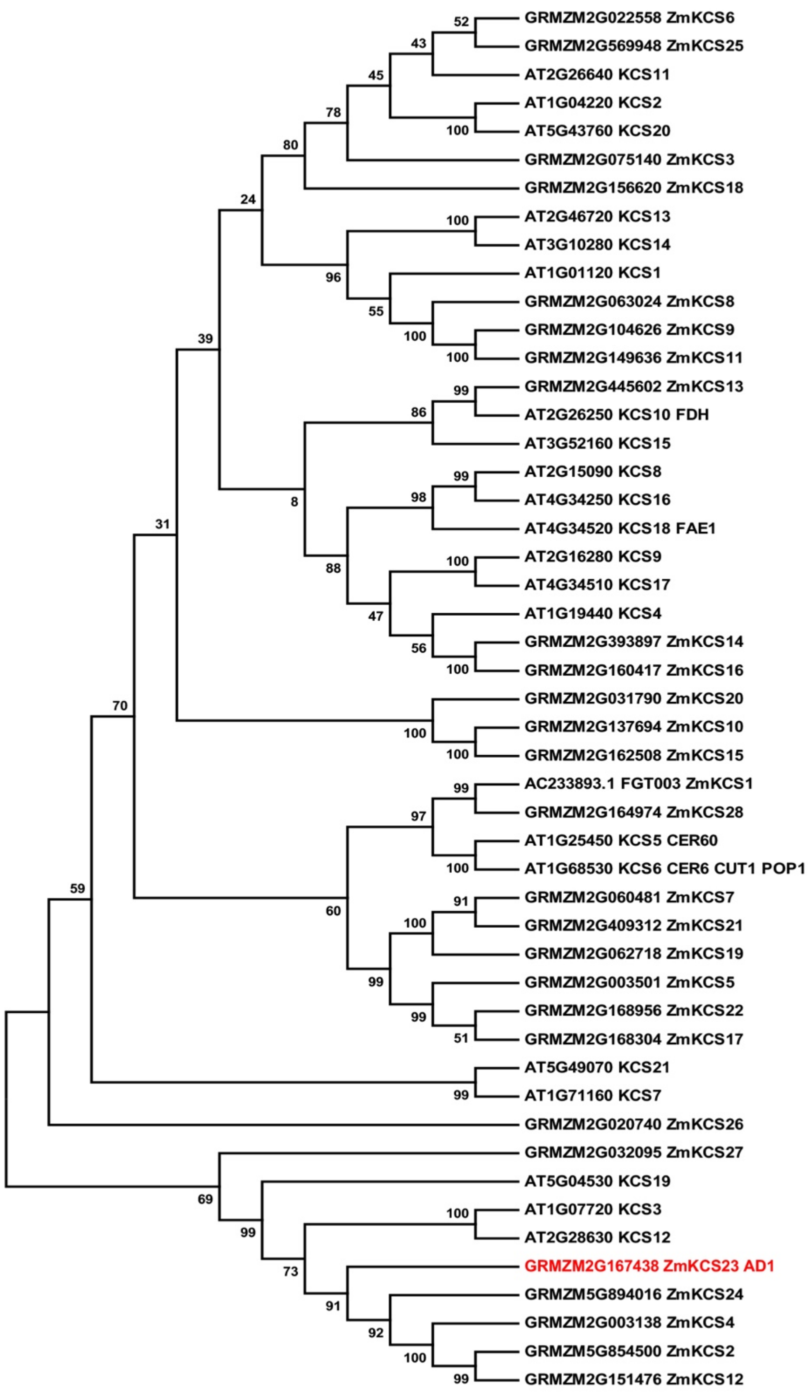


**Supplemental Figure 3.** Neighbor-joining phylogenetic tree of maize (GRMZM) and Arabidopsis (AT) KCS family proteins. Numbers represent reliability levels based on 1000 boostrap replications.

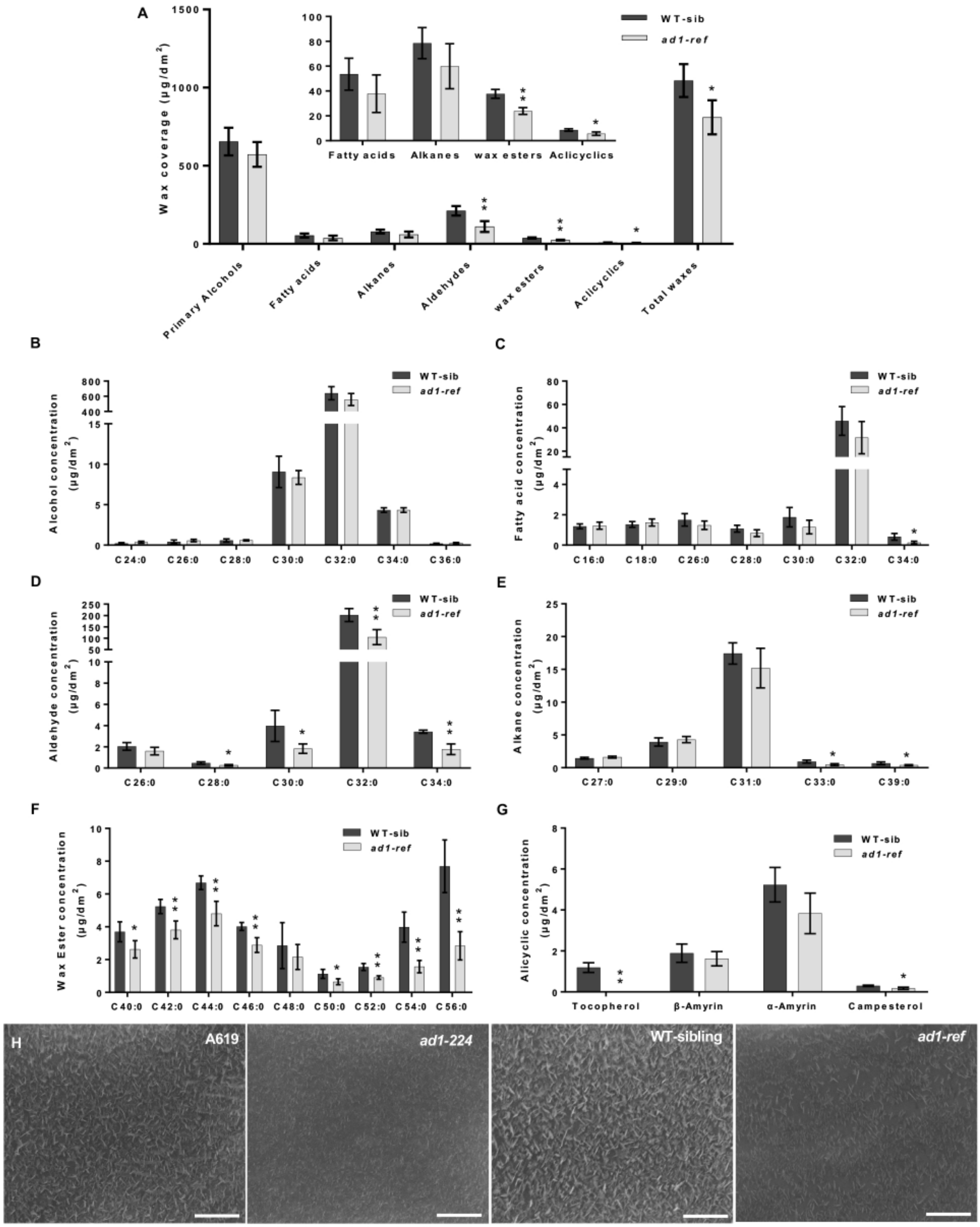


**Supplemental Figure 4.** Cuticular wax analysis in *ad1-ref* mutants.
(A) Total wax coverage and amount of each wax class in *ad1-ref* mutants and wild type leaves. The inset shows the less abundant wax classes at a different scale to more

clearly visualize significant differences. (B-G) Concentration of individual components in each wax class. primary alcohol (B), fatty acid (C), aldehyde (D), alkane (E), wax ester (F) and alicyclic (G). Means of 4 replicates and SD are reported. The asterisks represent significant difference determined by the Student’s test, * $P < 0.05$; ** $P < 0.01$. (H) SEM images of epicuticular wax crystals on the third leaves in A619, *ad1-224*, WT-sibling and *ad1-ref* samples used for the analysis. Scale bars, 5 μm.

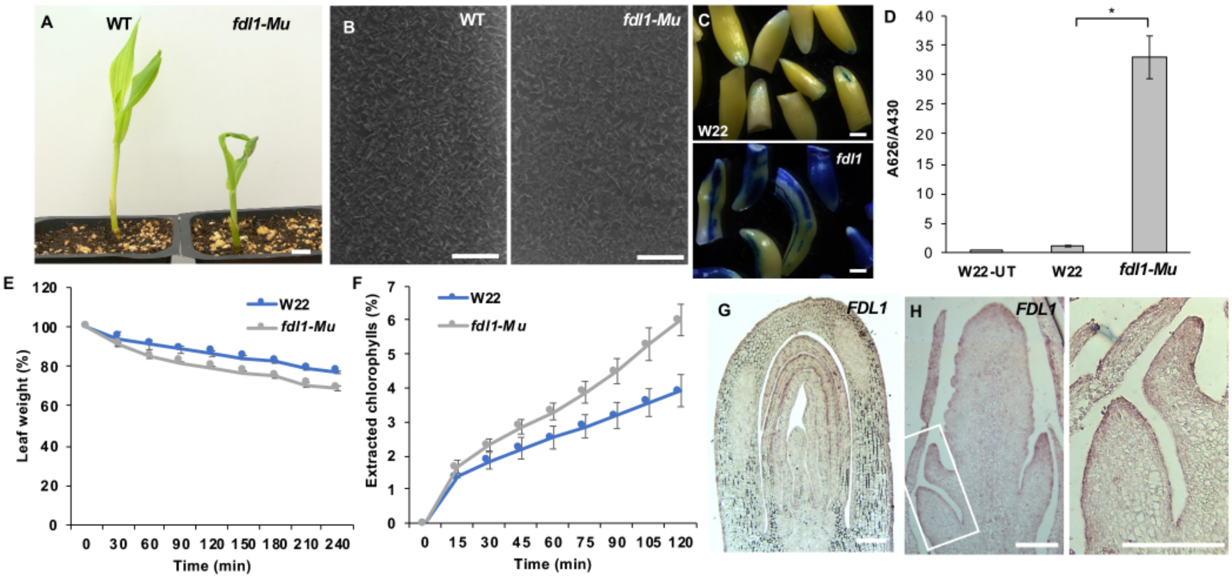


**Supplemental Figure 5.** Analysis of *FDL1* function.
(A) The *fdl1-Mu* mutant shows a leaf adherent phenotype at the seedling stage. (B) SEM images of epicuticular wax crystals on the third leaves in wild type and *fdl1-Mu*. Scale bars, 5 µm. (C) Toluidine blue tests on etiolated coleoptiles. Scale bars, 1 mm. (D) Quantification of toluidine blue uptake by the coleoptile of young seedlings, normalized to chlorophyll content. n=5 (5 seedlings per repetition; * Student-test P < 0.001). Error bars represent SD. (E) Water loss assays. Error bars show SD, n=3. (F) Chlorophyll leaching assays. Error bars show SD, n=3. (G,H) mRNA *in situ* hybridizations using *FDL1* antisense probes. Expression pattern in germinating seedlings (G) and tassel (H). In (H), right image is a higher magnification of the area framed in the left panel. Scale bars, 0.2mm.

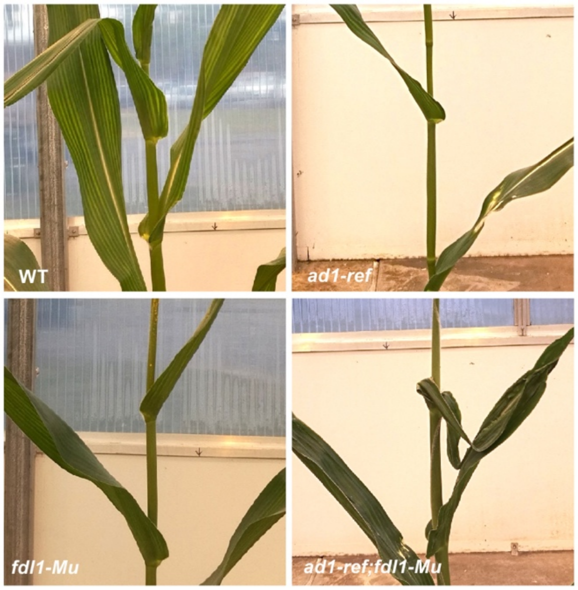


**Supplemental Figure 6.** Mature leaf fusion defects in single and double *ad1 fdl1* mutant adult plants.

**A**

GGGAAAACTATGGATAGACTGCTGCAGACGGCCCTAGAACTTTGACTAAACTTCAGTTGACTTGTTAGCATTTCTATTTTGTTTTTAGGGGC
TAAAGTTTAGTAGCTACTTGAGATCCAGATAGTTTTGTAAGAACTTTTTTCATTGTTGGGTTCATTACAATTTTTTGTAACGAAGTACAATT
CCGTGTTGCCCTAAATGCATTGCTGGTAGTCCTAGCACGCATGTTACTCGAATAGTATATAGCAAAGGACAAAAAAGAGGACAACATTTATG
GTCAAGAAAGTAAGACGATTTGTACAAAACTAAATATATGCTTATTTTTAGTCGTCGTCCTGAAGTATAGATGCTTAATGAGCCTCCTCCTC
CAGCGGCAGTGAAGTCATCCAATGCACAGTTATTAGAATCAGATGAAACATCAAACCAACGTTGCTGGTTCACGTGGTTACGGTTTGATTGA
GGATCCAACCAGCACTGTTAATGCTAAAGTAAAAAAATGGGCCATGCAAGAAATAGTAGATGAGGGAAGGTTCTCATGAGCCATGCATGCAT
GCATGATGACCCATGTCCATGATGTGGTATCCCCTTCTGCGTGCTATAAAACCACCCTCCACCCTCCTCCCTTTCCCCAACCTCTGAACCCC
GCTCTAATAAATCTTCCTCCCATCTCTCTCGCTCCCAACCCTCAAGTCCCCACCAACCCAACCCTGGTGGCACCGGCCGCACACACGAAAGC
AAGCTAGCTAGTCACTATTAGAGCTATTTGGTTGGGAGCGGTTTAGCTGAGCAAGATCGAACTGGCATTGTCACACACATCTCAGCACTCAG
CTCCTTGGGTTCCCCTCCATGAGCAGCAGCTGAGCAGCCACACATTGCCGTCTTCCTTGCTGCACTTGTAGTTACAGCTAGATCCATTGATT
TTTTTGGCAACAACAAGAGCAGGAGCTACTGAGCAAGAAGTGCGTACGTGCTTGGTGAGTGTTACTTATACTAGCTTGCGATG

**B**

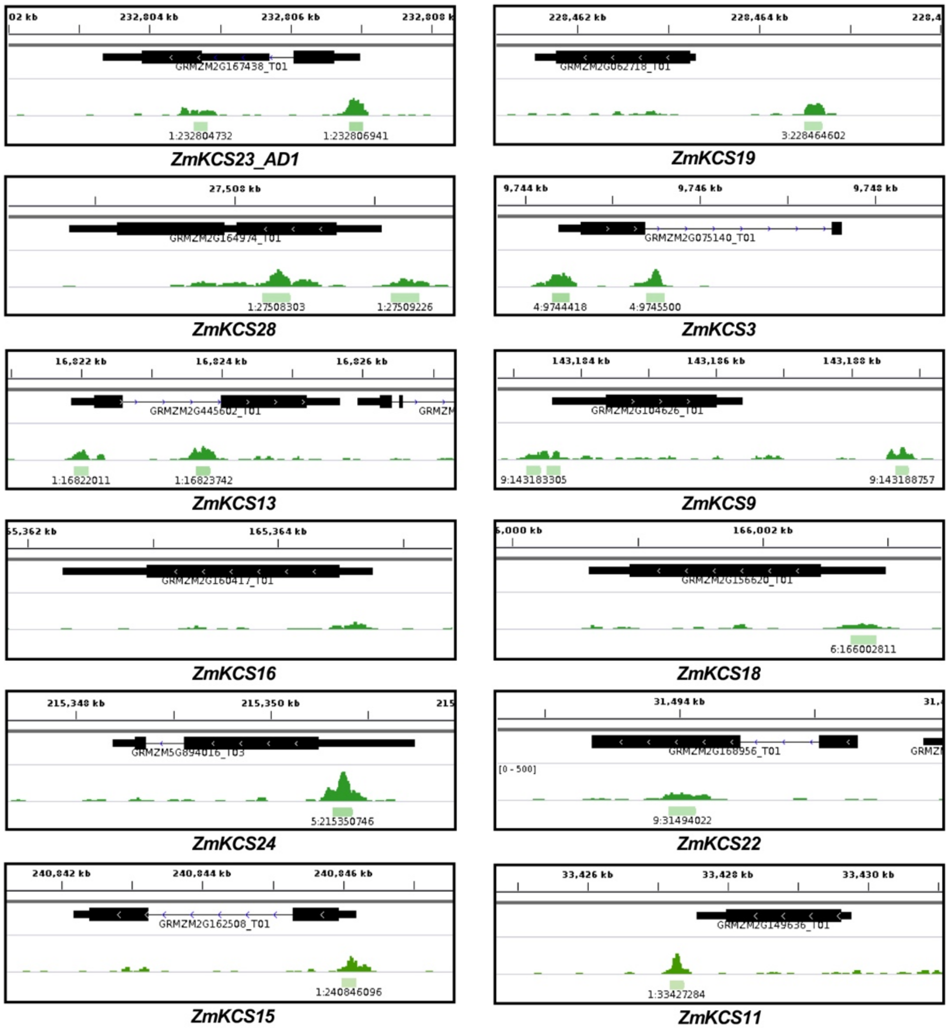


**Supplemental Figure 7.** FDL1 binds to the regulatory regions of several *KCS* genes. (A) *AD1* promoter sequence (-1000 to -1). Core binding motif marked in red. Gray background indicate 5' UTR region. (B) Genome browser screenshots of several *KCS* genes bound by FDL1.

**Supplemental Table 1.** List of wax biosynthetic genes directly bound by FDL1 in DAP-seq. * indicates gene with peak called only with default threshold. Gene IDs with no asterisk correspond to peaks with adjusted p-value > 1e-5. Gene IDs in bold are those whose close homologs were previously reported to be bound by AtMYB94 or AtMYB96.

| Gene ID | Annotation | Gene symbol |
|---|---|---|
| GRMZM5G858094* | acetyl-CoA carboxylase 2 | ACC2 |
| GRMZM2G062718* | 3-ketoacyl-CoA synthase | ZmKCS19 |
| **GRMZM2G075140** | **3-ketoacyl-CoA synthase** | **ZmKCS3** |
| **GRMZM2G104626*** | **3-ketoacyl-CoA synthase** | **ZmKCS9** |
| **GRMZM2G149636*** | **3-ketoacyl-CoA synthase** | **ZmKCS11** |
| **GRMZM2G156620*** | **3-ketoacyl-CoA synthase** | **ZmKCS18** |
| GRMZM2G160417* | 3-ketoacyl-CoA synthase | ZmKCS16 |
| GRMZM2G162508* | 3-ketoacyl-CoA synthase | ZmKCS15 |
| **GRMZM2G164974** | **3-ketoacyl-CoA synthase** | **ZmKCS28** |
| GRMZM2G167438 | 3-ketoacyl-CoA synthase | ZmKCS23 |
| GRMZM2G168956 | 3-ketoacyl-CoA synthase | ZmKCS22 |
| GRMZM2G445602* | 3-ketoacyl-CoA synthase | ZmKCS13 |
| GRMZM5G894016 | 3-ketoacyl-CoA synthase | ZmKCS24 |
| AC205703.4_FG006 | beta-ketoacyl reductase | KCR1 |
| **GRMZM2G075255** | **Fatty acid hydroxylase** | **CER1** |
| GRMZM2G029912 | Fatty acid hydroxylase | CER3 |
| GRMZM2G114642* | Fatty acid hydroxylase | CER3 |
| GRMZM2G104847* | long-chain acyl-CoA synthetase 2 | LACS2 |
| **GRMZM2G077375*** | **O-acyltransferase (WSD1-like) family protein** | **WSD1** |
| GRMZM2G083725 | glycosylphosphatidylinositol-anchored lipid protein transfer 1 | LTPG1 |
| AC197146.3_FG002* | myb domain protein 30 | MYB30 |
| GRMZM2G001223 | myb domain protein 106 | MYB106 |

**Supplemental Table 2.** List of primers used in this study.

| PRIMERS | SEQUENCE 5'-3' | PURPOSE |
|---|---|---|
| ad1-224 | For: TGTCAGTCATGGATGCTGGATTACG<br>Rev: CATGAAGGTGCGCTTGTTGCCCGGGTA | dCAPS KpnI - genotyping |
| ad1-ref | For: TGTCAGTCATGGATGCTGGATTACG<br>Rev: CTCAGCTAGCATTGTTGTGCTCACC | genotyping |
| fdl1-Mu | For: GAGTACACGAGCAATCCGCAA<br>Rev: CGCCAGGTTGATGTCCGTCT | genotyping |
| UBI-RT | For: GAGTGCCCCAACGCCGAGTG<br>Rev: CTACGCCTGCTGGTTGTAGACGTA | Real-time PCR |
| AD1-RT | For: TCTCCAACCCATTCATGGACAA<br>Rev: TTGAGACGTTGAGTTGAGCAGCT | Real-time PCR |
| ZmKCS13-RT | For: ATGGACGCCTAGCACGTACCT<br>Rev: GGTGAAGTGCAGCACTGCACT | Real-time PCR |
| ZmKCS15-RT | For: GAGACCACCTACAAGTTCGCTGA<br>Rev: GCTGATGATGAGAAATCAATCATGT | Real-time PCR |
| ZmKCS24-RT | For: GGAGGACTGCATCGACCAGTA<br>Rev: TCGTTAACCCATCCGAACTTGT | Real-time PCR |
| AD1-insitu-Probe | For: GAATTCGGCCGTCAAGGCCACCCTAGCTGCTCAACTCAACGT<br>Rev: AGTCGACGGCCCATGAGGCCCAGGACGACACTGACTGATTGACT | *In situ* |
| FDL1-insitu-Probe | For: GAATTCGGCCGTCAAGGCCAGTGTCCAATCGATCAAGCTCAA<br>Rev: AGTCGACGGCCCATGAGGCCACTTGCACAAGGGGATCTACGTA | *In situ* |
| AD1-EMSA-Probe | For: GACCCATGTCCATGATGTGGTAT<br>Rev: CCAAATAGCTCTAATAGTGACTA | EMSAs |
| AD1-Pro-mut1 | For: TTCAAGTCCCCATTTTTTTTTTTTTGGTGGCACCGGCCGCACAC<br>Rev: AAAAAAAATGGGGACTTGAAAAAAAAAAGCGAGAGAGATGGGAGG | EMSAs |
| AD1-Pro-mut2 | For: CCCTTTTTTTTTTTTCTGAACCCCGCTCTAATAAATCTTCCT<br>Rev: GGTTCAGAAAAAAAAAAAAGGGAGGAGGGTGGAGGGTGGTT | EMSAs |
| pGreenII-AD1Pro | For: TCGAGGTCGACGGTATCGATCCACACAATCGAGCACGACAT<br>Rev: CGGCCGCTCTAGAACTAGTGCGCAAGCTAGTATAAGTAACA | Transient Transcription Assay |
| pRI101-Myb94 | For: AAGTTCTTCACTGTTGATACATGGTGAGGCCGCCGTGCTG<br>Rev: AGTTGTTGATTCAGAATTCGCTAGAAGGGGTAATCCATGG | Transient Transcription Assay |
